# The curious case of cyanobacteria: a tale of light and darkness

**DOI:** 10.1101/2023.05.16.541008

**Authors:** Victoria Lee, Sally Zheng, Isaac Meza-Padilla, Jozef I. Nissimov

**Author notes:** equal contribution (Victoria Lee, Sally Zheng). Competing interests: The authors declare no competing interests.

## Abstract

Toxic algal bloom-forming cyanobacteria are a persistent problem globally for many aquatic environments. Their occurrence is attributed to eutrophication and rising temperatures due to climate change. The result of these blooms is often loss in biodiversity, economic impacts on tourism and fisheries, and risks to human and animal health. Of particular concern is the poorly understood interplay between viruses and toxic species that form blooms because viruses may exacerbate their harmful effects. Concurrently, cyanobacteria are also a source of bioactive compounds other than toxins, which makes them good candidates for drug discovery. We show that virus infection of the cyanobacterium *Microcystis aeruginosa*, results in as high as a 40-fold increase in the toxin microcystin two days post virus infection (dpi), and predict that microcystin levels may remain high in a body of water up to 7 dpi, long after water discoloration and cell lysis. This implicates viruses as major contributors to toxin release from cyanobacteria and emphasizes the importance of taking them into account in predictive models and in the assessment of water safety. We also show that bioactive compounds of *M. aeruginosa* inhibit and delay infection of single stranded DNA and single stranded RNA viruses. This highlights the potential of cyanobacteria as an excellent source for the discovery of novel antiviral compounds, and the ease with which screening for cyanobacterial antivirals can be achieved.

## INTRODUCTION

*Microcystis aeruginosa* (*M. aeruginosa*), a globally distributed freshwater harmful algal bloom (HAB)-forming cyanobacterium, produces a variety of secondary metabolites that are not directly associated with growth, development and reproduction of the cell (*Humbert et al., 2013*). While some of these are bioactive compounds such as the liver damaging hepatoxin microcystin, and the neurotoxic cyanopeptolins that affect the nervous system (*Mayer et al., 2011*), other ones include fatty acids, vitamins C and E, antibacterials, and antivirals (*Gupta et al., 2013*). Therefore, it can be stated that the physiology of *M. aeruginosa* has both negative and positive aspects.

Recent studies hint at the fact that the harmful effects of toxin producing species of cyanobacteria may be exacerbated by cyanophage infection (i.e., viruses that infect cyanobacteria), where cell lysis triggers the excess release of toxins from the infected cells (*Šulčius et al., 2018; McKindles et al., 2020*). On the other hand, studies show that crude *M. aeruginosa* extracts inhibit influenza A in cell culture (*Nowotny et al., 1997*), and that a recently isolated lectin from this bacterium, microvirin, inhibits HIV-1, also in cell cultures (*Kehr et al., 2006; Huskens et al., 2010)*, which hints at the beneficial effects of *M. aeruginosa*. Nevertheless, the degree to which cyanophages affect the production and release of toxins by HAB-forming cyanobacteria, and by extension, the implications this may have on water safety, is poorly studied, as is the ability of cyanobacterial antivirals to inhibit other types of viruses.

Therefore, using *M. aeruginosa* strain NIES-298 and its cyanophage Ma-LMM01 (Table S1) as an experimental model system, we first performed infection experiments to characterise the temporal dynamics of microcystin during virus infection. We were able to empirically show elevated levels of microcystin in infected treatments after cell lysis and predicted the time it would take post virus infection for the toxin’s concentration to reach acceptable levels of 1.0 ppb (Figure 1); the World Health Organization (WHO) recommended upper limit of microcystin-LR in drinking water (*WHO, 2020*). Then, using our extensive isolate collection of marine (e.g., diatoms and coccolithopores) and freshwater (e.g., chlorella) microalgae and their viruses (Table S1), we also embarked on a proof of concept study, in which we tested the inhibitory activity of filtrates from three *M. aeruginosa* strains (e.g., NIES-298, CPCC 299, and CPCC 300) against a diverse range of non-pathogenic naked ssDNA (e.g., CtenDNAV), ssRNA (e.g., CtenRNAV) and enveloped dsDNA viruses (e.g., *Emilianiaa huxleyi* virus 201 and 99B1; and chloroviruses IL-5-2s1 and NY-2A). We showed that inoculating virus-infected samples with late exponential growth phase filtrates from the three cyanobacterial strains delayed infection by some of the viruses we tested (Figure 2), suggestive of the presence of antiviral compounds.

**Figure 1.**
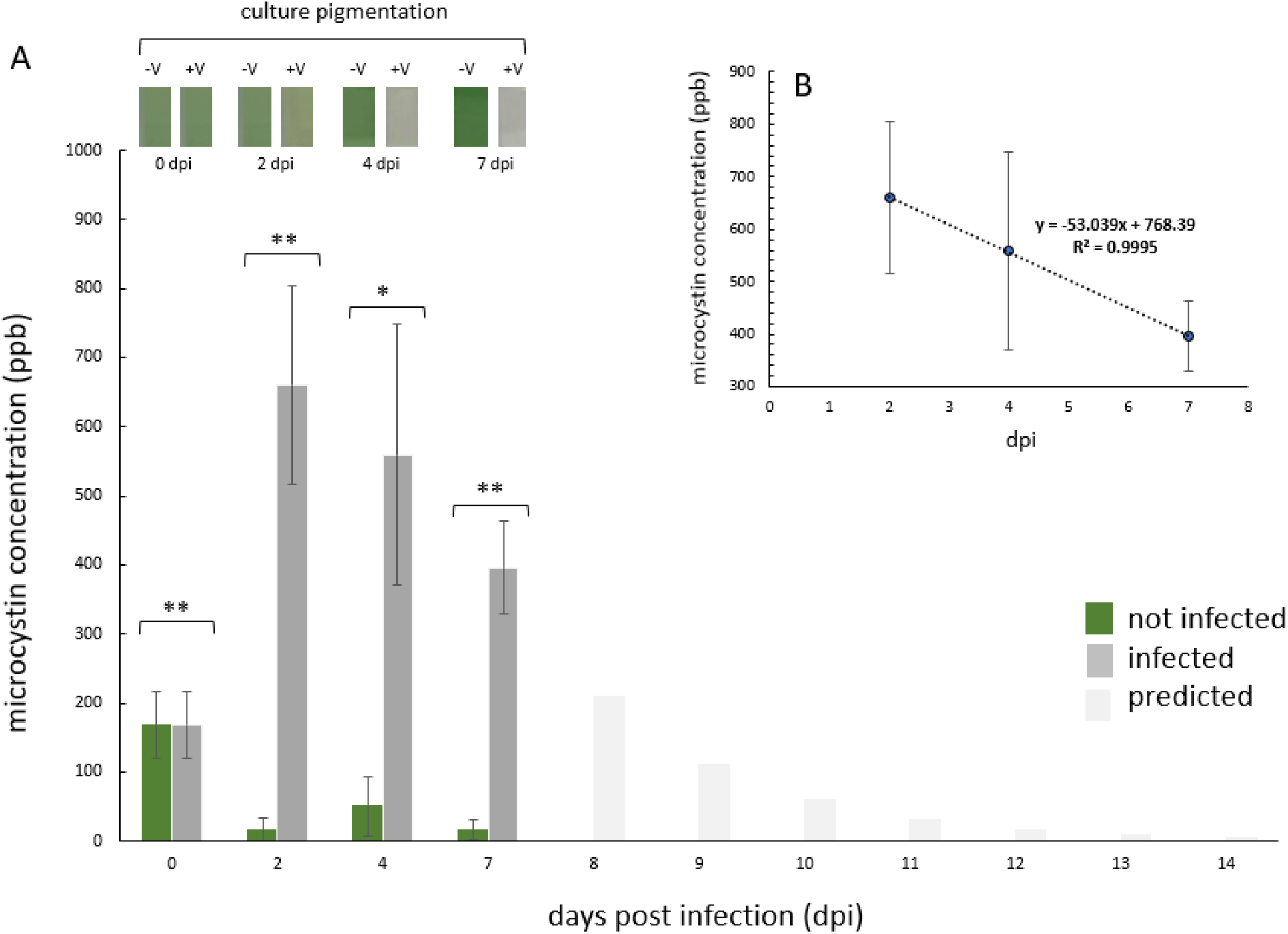
Microcystin dynamics during cyanophage infection of *M. aeruginosa*. (**A**) Average (± SD, *n* = 3) microcystin concentration in parts per billion (ppb) in cyanophage Ma-LMM01 infected (dark grey bars) and non-infected (green bars) treatments. ** and * denote significant differences (p<0.01 and p<0.05 respectively; ANOVA). Predicted microcystin concentrations (light grey bars) are shown for 8-14 dpi, based on the calculated daily average decrease in microcystin concentration throughout days 2-7. Inset represent culture pigmentation (photographed and cropped) of a representative triplicate treatment with either viruses (+V) or without (-V) 0-7 dpi. (**B**) Average daily rate of microcystin decrease in infected treatments, calculated between the highest measured concentration on day 2, until the last day of the experiment on day 7.

**Figure 2.**
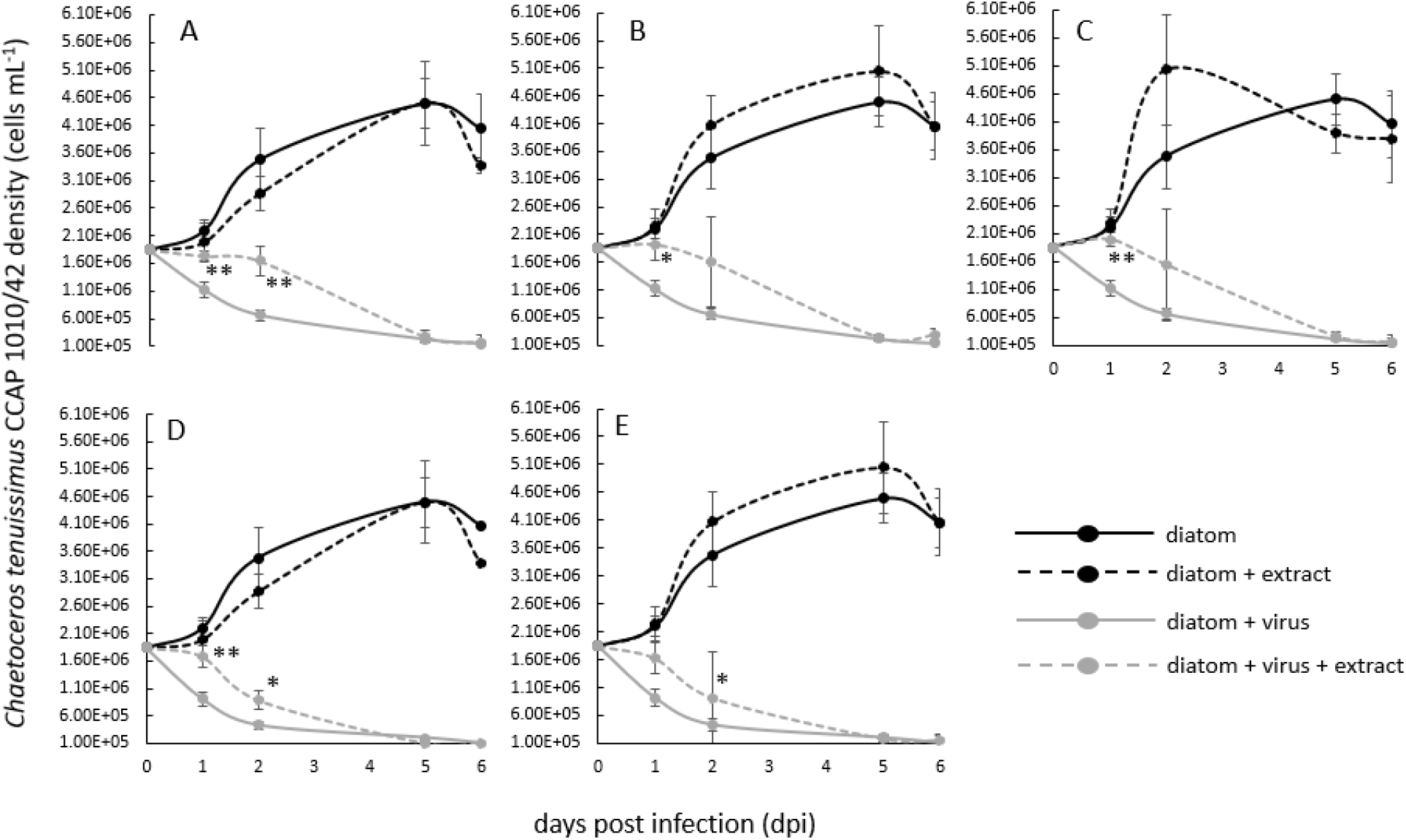
*Chaetoceros tenuissimus* strain 1010/42 growth dynamics in experiments that tested the effects of late exponential growth phase extracts (i.e., filtrates) from *M. aeruginosa* NIES 298 (**A & D**), CPCC 299 (**B & E**) and CPCC 300 (**C**), on CtenDNAV (**A, B**, & **C**) and CtenRNAV (**D & E**). Solid black lines= diatom cells only; dotted black lines= diatom cells with added extract; solid grey lines= diatom cells without extract but infected with CtenDNAV or CtenRNAV; dotted grey lines= diatom cells with added extract and also infected with CtenDNAV or CtenRNAV). ** and * denote significant differences (p<0.01 and p<0.05 respectively; ANOVA) between cultures (± SD, *n* = 3) infected by only a virus and those that were infected with viruses but also contained added cyanobacterial extracts. Note that no inhibition of chloroviruses and coccolithoviruses was observed by any of the extracts tested. Inhibition of CtenRNAV by CPCC300 extracts was also not observed, hence data for these is not shown.

Collectively our study shows that while cyanobacterial bioactive compounds such as toxins are induced by viruses to extremely high levels, levels that have clear ecological and human health implications, they may also have an unexplored as a of yet applied potential, and are worthy of further study.

## RESULTS AND DISCUSSION

### Tale One-‘The dark side’ of toxic cyanobacteria

The first part of our study involved analysis of total microcystin concentration during virus infection (Figure 1 & S1; Table S1). The starting average (n=3) microcystin concentration (ppb) in our cultures prior to virus infection was 167.76 ppb (±48.47). After infecting triplicate cultures of *M. aeruginosa* NIES-298 with Ma-LMM01 (Figure S1), the concentration of microcystin remained extremely high throughout the experiment compared to the uninfected controls (Figure 1-A). The highest concentration (∼660.34 ppb, ± 144) was measured two days post infection (dpi), where it was an average of ∼40-fold higher than in the respective non infected treatments, and represents a concentration increase of nearly 4-fold in two days (i.e., total average increase of 492.58 ppm, ±96.55). This initial increase is a result of the lysis of 2.49 × 10^7^ cells mL^-1^ (± 5.5 × 10^5^ cells mL^-1^), measured here as a decrease in cell abundance from T-0 to 2 dpi (Figure S1-B).

Assuming an equal microcystin release for each lysing cell due to infection, this means that virus induced cell lysis was responsible for the release of 1.98 × 10^−5^ ppb cell^-1^ in the first two days. Notably, natural *M. aeruginosa* abundances are much lower than in our laboratory experiments. However, if we are to extrapolate our calculated per cell microcystin release due to virus infection to a natural system, then a complete lysis event in a location such as Lake Agawam, in which *M. aeruginosa* abundances can be as high as 1.48 × 10^6^ cells mL^-1^ (*Davis et al., 2009*), will result in a total increase of ∼29 ppb. This is much higher than the provisional recommended upper limit for drinking water by the World Health Organization (WHO), which for the most common and toxic microcystin, microcystin-LR, is 1.0 ppb (*WHO, 2020*). And although the average microcystin concentration remained high until the end of our experiments on day 7, it appeared to decrease daily by an average of 0.53-fold (Figure 1-B). Based on this daily decrease, it can be calculated that it may take >14 days post infection until the levels of this toxin drops to below the recommended level by the WHO in our experiments (Figure 1-A), or in the case of a natural setting such as Lake Agawam, >7 days post infection/the onset of host cell lysis.

It is important to note that the real numbers in a natural, and more complex system, might be different than those provided in the crude calculations here, because they do not take into account other physicochemical and biological factors that are at play during host-virus-toxin dynamics in an *in situ* algal bloom (e.g., temperature, pH, nutrient loads and trophic status of a system, presence/absence of grazers, host resistance to infection, diversity and abundance of other microorganisms and viruses, virus decay rates, bio- and photo-degradation of toxin, etc.). Also, our total microcystin measurements did not follow the traditional protocol that includes mechanical lysis of cells, because this would have made it difficult to differentiate between virus-induced toxin increase and increase due to mechanical lysis. This means that our numbers are likely an underestimate. However, to the best of our knowledge, direct empirical measurements and quantification of virus induced toxin release from infected *M. aeruginosa* cells in a controlled setting have not been reported yet. Hence, this work is an important “springboard” for the inclusion of viruses in HAB dynamics modelling, as predictive models rely on the responses of toxic species to both abiotic and biotic drivers (*Litchman et al., 2023*).

Further, the fact that high levels of microcystin are still being measured in parallel to water discoloration (inset of Figure 1-A) and cyanobacterial abundance decrease due to virus infection (Figure S1-B), is notable, and reinforces the importance of actively measuring toxin levels whenever possible, rather than relying solely on cyanobacterial biomass and chlorophyll measurements, or the quantification of genes involved in toxin production. Indeed, water quality guidelines reiterate that water monitoring should be a multifaceted approach, including the use of enzyme–linked immunosorbent assays (ELISA), liquid chromatographic methods, and protein phosphatase inhibition assays (Health Canada, 2022). Due to the expense of these assays and the need for specialised infrastructure/trained personnel, these more advanced approaches may not always be an affordable option, especially for those in low income or developing countries. The use of mobile apps (e.g., the Cyanobacteria Assessment Network Application app developed by the United Stares Environmental Protection Agency, and the Bloomin’ Algae app developed by the UK Centre for Ecology and Hydrology, to name a few) that relay on satellite data to detect and map the location of cyanobacterial blooms can help decision makers in rapidly detecting potential HABs. However, given that they are based on water discoloration, they too are less useful in situations that are characterised by high viral activity and low cyanobacterial biomass, lacking the ability to accommodate for virus-induced toxin release from cells and subsequent toxin proliferation over time.

### Tale Two-‘The bright side’ of toxic cyanobacteria

In the second part of our work, we spiked with *M. aeruginosa* NIES-298, CPCC 299, and CPCC 300 “extracts” (i.e., 0.45 μm filtrates that excluded cells), virus-infected cultures of *Chlorella variabilis* NC64A, *Emiliania huxleyi* CCMP374, and *Chaetoceros tenuissimus* CCAP1010/42 (e.g., infected by dsDNA chloroviruses IL-5-2s1 and NY-2A; dsDNA coccolithoviruses EhV-201 and EhV-99B1; and ssDNA CtenDNAV and ssRNA CtenRNAV diatom viruses, respectively) in order to explore the antiviral potential of the bioactive compounds released by cyanobacteria (Figure S2 & S3). The extracts showed no inhibitory effects against the coccolithovirus and chlorovirus strains (data not shown). However, extracts from all three cyanobacterial strains had an inhibitory effect against the diatom viruses CtenDNAV and CtenRNAV, as we observed delayed cell lysis up to two days post infection (dpi) in their presence (Figure 2).

Given that we did not quantify diatom virus production rates post infection or identified chemically the antivirals themselves, and only used the decline in host cell abundance as a proxy for successful virus infection/virus inhibition, we can only hypothesise at this stage about the mode of operation of these antivirals and their identity. Of the identified so far bioactive compounds that *M. aeruginosa* produces, only lectins and micropeptins have been documented to exhibit antiviral properties (*Mazur-Marzec et al., 2021*). The fact that we only observed inhibition of naked ssDNA and ssRNA viruses and not of the enveloped dsDNA viruses, points against the involvement of a lectin such as microvirin. This is because lectins tend to bind to the mannose-rich proteins of viral envelopes (*Mitchell et al., 2017*). Another possibility of course is that if the antivirals were indeed lectins, they simply may not have been effective against the dsDNA viruses we tested.

On the other hand, the fact that we only observed a delayed infection in the first two days, points towards inhibitory agents that operate in a dose dependent manner, such as micropeptins (or another unidentified in cyanobacteria as of yet antiviral). Micropeptins are serine protease inhibitors (*Thuan et al., 2019*) that can affect virion packaging by blocking access to host proteases, necessary for allowing a virus to complete its infection cycle (*Steinmetzer and Hardes, 2017*). Therefore, it is possible that if the antivirals were indeed micropeptines, then the added concentration in our experiment was not sufficient to completely block virion packaging and subsequent release, only delaying infection. Interestingly, extracts from all three *M. aeruginosa* strains show antiviral properties, suggesting that pharmacologically relevant compounds are an integral component of cyanobacteria more generally.

It was previously reported that antiviral defence genes are expressed by *M. aeruginosa in situ* (*Morimoto et al., 2019*) and an abundance of ssRNA viruses that infect photosynthetic protists such as diatoms seems to be present at the peak of Microcystis blooms (*Pound et al*.,*2020*). It was suggested that the large presence of non-cyanobacterial viruses during a HAB may help in eliminating competitors of *M. aeruginosa* by driving a system from a diatom dominated bloom in the winter, to a cyanobacterial dominance in the summer. It is tempting to hypothesize then a scenario in which the cyanobacterial antivirals act as an indirect “cyanobacterial suicide switch”, by targeting diatom viruses at the end of a summer and the autumn months, lifting diatom virus predation pressure, and allowing diatoms to become masters once again (Figure 3).

**Figure 3.**
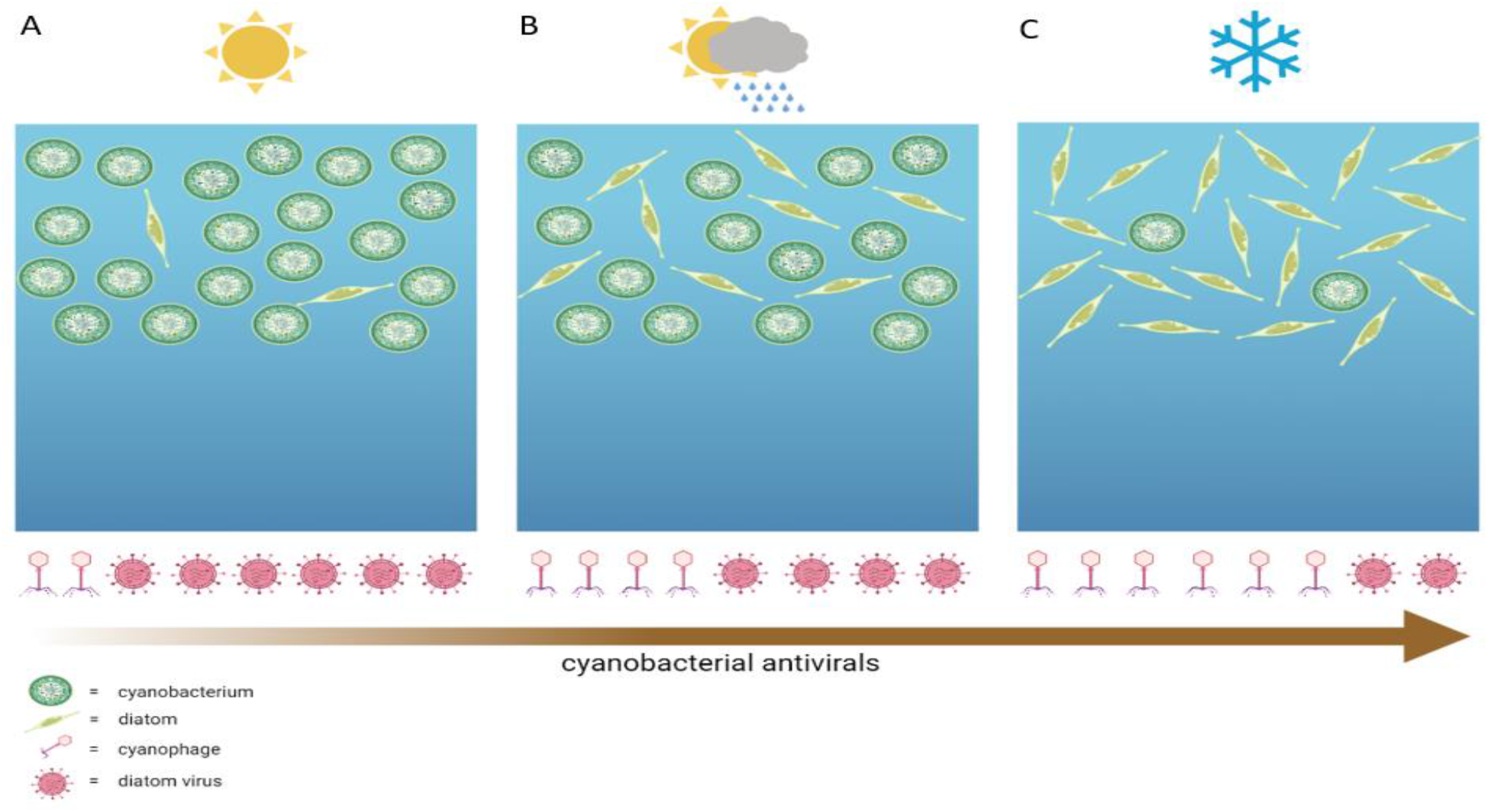
Hypothesized scenario of a temperate lake in which cyanobacterial antivirals act against diatom viruses and allow diatoms to proliferate in the winter. (**A**) Spring and summer months are typically characterized by high cyanobacterial biomass that can potentially cause HABs, in which initially the abundance and activity of cyanophages might be relatively low. (**B**) Cyanophages then increase in abundance and together with other biotic and abiotic factors help regulate the abundance of cyanobacteria, in the process of which there is an increase in the release of diatom specific antivirals from lysing cyanobacterial cells. (**C**) The cyanobacterial antivirals then act upon ssDNA and ssRNA diatom viruses, lifting predation pressure from diatoms and allowing them to become dominant again during the colder months. Note that cells and viruses are not drawn to scale and are for illustrative purposes only. Created in Biorender.

## CONCLUSIONS

To the best of our knowledge, this is the first time that high levels of microcystin are specifically being reported to occur long after cyanobacterial host lysis and loss in water coloration, with potentially real-world implications to lake habitats. This work re-emphasises the risk of ignoring the role of virus infection of HAB-formers; current HAB-monitoring guidelines lack a specific reference to the impacts of viruses. Further work should focus on studying in detail cyanoHAB-virus-toxin dynamics *in situ* over large spatial and temporal scales. At the same time, our work shines a positive light on toxic cyanobacteria, by implicating them as an invaluable resource for the discovery of novel antivirals. Although, the antiviral effects we observed were only in select few *M. aeruginosa* strains we had access to, there are hundreds of cyanobacterial species scattered in culture collections globally (Campbell and Lorenz, 2022), many of which may too prove to be good candidates for antiviral discovery. Finally, we demonstrated the ease with which cyanobacteria can be screened for antiviral activity, especially if testing these against already available in culture non-pathogenic viruses. Further work should assess the effects of different cyanobacterial extract concentrations, in parallel to: i) SEM and TEM imaging to reveal whether virus inhibition is at the point of entry, assembly, or exit from the infected diatom host cells, and ii) complete metabolomics analysis of the cyanobacterial extracts to elucidate chemically the antivirals that are at play.

## MATERIALS AND METHODS

### Microalgal and cyanobacterial growth conditions

Culture stocks of *M. aeruginosa* CPCC 299 and CPCC 300 cyanobacteria were obtained from the Canadian Phycology Culture Centre (CPCC) in Waterloo, Canada, whereas *M. aeruginosa* NIES 298 was obtained from the National Institute for Environmental Studies in Japan. All cyanobacterial strains were cultivated in BG-11 liquid media, at a light:dark cycle of 12:12 hours, and maintained at a temperature of 25°C (±1°C).

Culture stocks of the freshwater microalga *Chlorella variabilis* NC64A and the marine diatom *Chaetoceros tenuissimus* CCAP1010/42 were obtained from the Culture Collection of Algae and Protozoa (CCAP) in Scotland, and cultivated in Bold-Basal Medium with 3-fold nitrogen (3N-BBM) at 25°C (±1°C) and artificial seawater f/2-Si at 18°C (±1°C), respectively, at a light:dark cycle of 12:12 hours.

Culture stock of the marine microalga *Emiliania huxelyi* CCMP374 was obtained from our private culture collection at the University of Waterloo (Canada) and cultivated in artificial seawater f/2-Si at 18°C (±1°C), at a light:dark cycle of 12:12 hours.

The aforementioned culture conditions for each strain also apply to the ‘Tale 1’ and ‘Tale 2’ experiments outlined below.

### Virus strains

Fresh viral lysates of the virus isolates used in this study (Table S1) were obtained by infecting an exponentially growing host strain, filtering the lysed cells through a 0.45 μm filter, and storing the filtrates in the dark at 4°C until the onset of the experiments (Figure S3).

### Infectious titre quantification of virus lysates

To ensure that the virus lysates (Table S1) that we used in our experiments contained infectious particles, we conducted most probable number (MPN) assays (Suttle, 1993) on each lysate. Briefly, *M. aeruginosa* NIES 298, *Chlorella variabilis* NC64A, *Emiliania huxleyi* CCMP 374, and *Chaetoceros tenuissimus* 1010/42 cultures were grown to mid-exponential phase, after which 240 μL of each were distributed into individual 96-well plates. Then, a 10-fold serial dilution of each viral lysate was prepared. Subsequently, 10 μL of 0.2 μm-filtered media as a negative control were loaded in each well in column 1 of the 96-well plate (A1-H1), 10 μL of undiluted lysate were loaded as positive controls in each well in column 2 (B2-H2), and 10 μL of dilutions 10^−1^-10^−10^ were loaded in columns 3-12. The plates and the wells within them were visually inspected for infection (i.e., culture discoloration/loss in pigmentation relative to controls) up to 8 days post virus addition (i.e., culture discoloration in relation to controls). The scores (i.e., number of cleared wells for each dilution) were then inputted into the Environmental Protection Agency MPN Calculator to obtain an MPN number (i.e., the infectious number of virus particles in each lysate).

### Cell density measurements

Throughout our experiments we quantified host cell abundances with a haemocytometer (0.1 mm deep) under an optical microscope (Olympus BH-2) at the 40X magnification. For each sample we loaded 10 μL and counted the number of cells in 5 squares, from which we derived an average. While microalgal cells were placed onto the haemocytometer chamber directly, cyanobacterial counts were obtained after first heating up the cells at 60°C for 20 seconds. This was done because *M. aeruginosa* cells have gas vacuoles, which make them float in the counting chamber/hemocytometer grid and therefore difficult to count. The heating step bursts the vacuole and therefore allows cells to stay on the same plane of view while counting.

Error bars in the cell abundance graphs (Figure 2) represent ± 1 standard deviation (SD) of the mean of a given treatment, set up in triplicates (n=3). Statistical tests used a one-way analysis of variance (ANOVA) with a significance threshold defined by a p value of < 0.05.

### Infection experiments and toxin concentration measurements (Tale 1)

A 300 mL *M. aeruginosa* NIES 298 master culture was incubated until it reached a cell density of 2.65 × 10^7^ cells mL^-1^. This culture was then split into six 40 mL cultures (Figure S1-A). Three of the six cultures were infected with 2.5 mL of the Ma-LMM01 cyanophage lysate (which contained 1.35 × 10^7^ infectious viruses mL^-1^). The other three served as no virus control cultures, to which we added 2.5 mL 0.02 μm Ma-LMM01 cyanophage filtrate. Total microcystin (Adda Specific) concentration (i.e., cell associated and extracellular) measurements were then obtained with a commercially available enzyme–linked immunosorbent assay (ELISA) kit (Enzo Life Sciences) up to 7 days post infection (e.g., time of infection, 48h, 96h, and 168h) in accordance with the manufacturer’s manual, using a SpectraMax ABS Plus microtiter plate reader. In parallel, we obtained *M. aeruginosa* NIES 298 cell density counts (Figure S1-B) as previously described.

Error bars in the microcystin plot (Figure 1-A) represent ± 1 standard deviation (SD) of the mean of a given treatment, set up in biological triplicates (n=3). Statistical tests used a one-way analysis of variance (ANOVA) with a significance threshold defined by a p value of < 0.05.

### Calculated cellular microcystin release per cell and predicted degradation rate

To calculate the concentration of microcystin released per each lysed cell in our experiments (1.98 × 10^−5^ ppb), we first calculated the total average increase in microcystin in the first two days (492.58 ppb) post virus infection. We then divided this number by the total average number of cell decrease (cell loss) in that same time period (2.49 × 10^7^ cells mL^-1^). This allowed us to hypothesize the microcystin concentration increase in an *in situ* environment such as Lake Agawam, in which *M. aeruginosa* abundances were in August of 2006, 1.48 × 10^6^ cells mL^-1^ (*Davis et al., 2009*), if all the cells in that bloom were to be lysed (i.e., 1.48 × 10^6^ cells mL^-1^ × 1.98 × 10^−5^ ppb per lysed cell = 29.43 ppb).

To roughly calculate the daily rate of microcystin decrease in our experiments throughout the 7 days, we first averaged the measured fold decrease from day 2 to day 4 (660.34 ppb/559.50 ppb/2 days= 0.59), and from day 4 to day 7 (559.50 ppb/395.8 ppb/3 days= 0.47), and then averaged the two numbers, which resulted in an average daily fold decrease of ∼0.53. Assuming a similar daily fold decrease in measured microcystin beyond day 7, it can be calculated that at day 14 post virus infection, the microcystin levels would be 4.6 ppb, and at day 15, it would be at undetectable levels. Assuming a similar daily fold-decrease in a natural setting such as Lake Agawam, where we estimated that microcystin concentration 2 dpi to be 29.43 ppb, it means that it would take >7 days for the levels to reach below 1 ppb (29.43 ppb*0.53= 15.6 ppb at day 3; 15.6 ppb*0.53=8.27 at day 4; and so forth). Note that these calculations do not take into account the volume in which the microcystin would become dissolved and hence its half-life.

### Infection experiments evaluating the antiviral activity of cyanobacterial extracts (Tale 2)

Before testing the antiviral effects of bioactive compounds from cyanobacteria, we obtained cell-free extracts (filtrates that excluded cells) from exponentially growing *M. aeruginosa* NIES 298, CPCC 299, and CPCC 300 (Figure S3-A). To do so we first cultivated 100 mL of each strain. Then, 15 mL from each strain were 0.45 μm syringe filtered at the late exponential growth phase. The filtrates, termed hereon the cyanobacterial extracts, were then stored at 4°C until the beginning of our spike experiments (Figure S3-A).

For each spike experiment (Figure S3-B), master cultures of *E. huxleyi* CCMP374, *C. variabilis* NC64A and *C. tenuissimus* CCAP1010/42 were cultivated until mid-exponential phase, after which 1.6 mL aliquots were dispensed into 24-well microculture plates as per the outline in Table S2. For each infection experiment with the aforementioned algal hosts and their respective virus isolates (see Table S1) we included a control plate with only the algae (1.6 mL), no added viruses, and an added 200 μL of BG-11 media, to a total of 1.8 mL per each well (e.g., control plate 1); a control plate with the algae (1.6 mL), no added virus, added 100 μL of BG-11, and added 100 μL of a cyanobacterial extract, to a total of 1.8 mL in each well (e.g., control plate 2); a control plate with the algae (1.6 mL), added 100 μL of BG-11, and added 100 μL of virus, to a total of 1.8 mL in each well (e.g., control plate 3); and an experimental plate that consisted of 1.6 mL of algae, 100 μL of a cyanobacterial extract, and 100 μL of virus. The plates were then incubated at their standard optimal growth conditions (see above) up to 6 days, with cell abundance measurements taken as previously described at day 0, 1, 2, 5 and 6.

Note that the reason we chose to supplement some of the wells with BG-11 media is because: 1) we wanted to keep a consistent total volume of 1.8 mL across wells and treatments, 2) BG-11 would not have affected the growth dynamics of *E. huxleyi, C. variabilis* or *C. tenuissimus*, and 3) this media is also a constituent of the cyanobacterial extracts that were spiked in each experiment.

## ACKNOWLEDGEMENTS

We thank Dr. Yuji Tomaru from the Fisheries Technology Institute in Japan for allowing us to use their diatom and cyanophage virus isolates; and to Dr. David D. Dunigan from the Nebraska Center for Virology for allowing us to use their chlorovirus isolates. We also thank the National Institute for Environmental Studies (NIES) in Japan, the Canadian Phycological Culture Centre (CPCC), and the Culture Collection of Algae and Protozoa (CCAP) in Scotland, for allowing us to purchase some of the microalgal and cyanobacterial strains used in our study. This work was funded by an NSERC Discovery Grant (2022-03350) and a Phycological Society of America Fellowship awarded to JIN.

## AUTHOR CONTRIBITIONS

VL, SZ, and JIN designed the study. VL conducted infection experiments, counted cells, and quantified microcystin concentrations (Tale1); SZ conducted infection experiments, counted cells, and tested extracts for antiviral activity (Tale 2); and IMP conducted MPN assays to quantify the infectious titre of the viruses used in this study. VL, SZ, and JIN wrote the manuscript. All authors edited and approved the final version of the manuscript.

## ADDITIONAL FILES

Supplementary Figures File (Figures S1-S3; Tables S1 & S2).

## Notes

### Competing Interest Statement

The authors have declared no competing interest.

