## SUPPLEMENTARY MATERIALS for "The curious case of cyanobacteria: a tale of light and darkness"

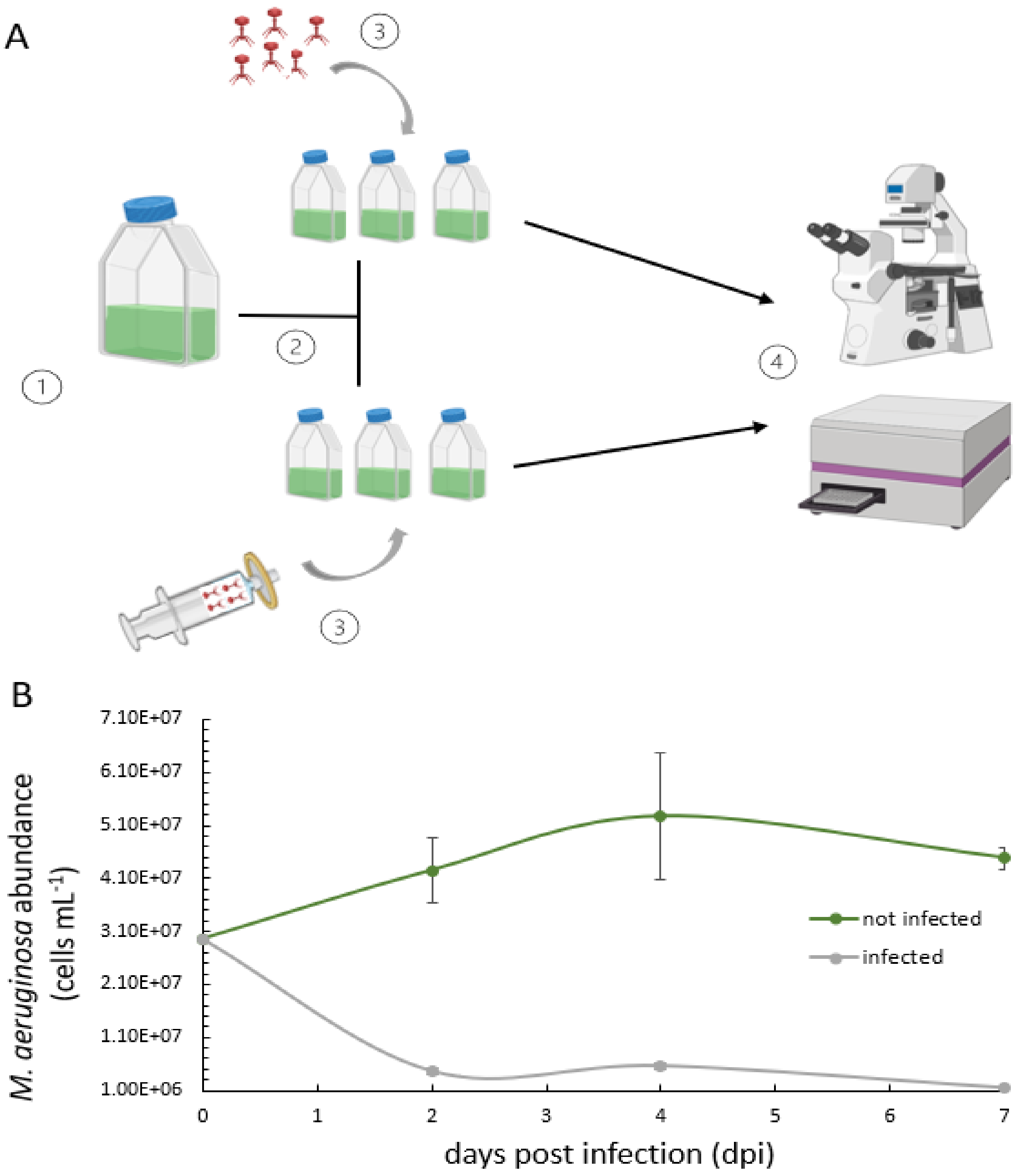


**Figure S1.** Experimental set up of cyanophage infected and non infected *M. aeruginosa* NIES 298 cultures and their subsequent analysis. (**A**) 1- Cyanobacterial cells were incubated in a master culture until mid-exponential phase; 2- the culture was then split into six replicates, three of which were infected with a cyanophage Ma-LMM01 stock that was at a virus particle density of 1.35 × $10^7$ $mL^{-1}$, and three inoculated with an equal volume of 0.02 µm filtrate of the Ma-LMM01 stock; 4- ELISA essays and total cell abundance measurements, were performed using spectrophotometry and haemocytometery, respectively. (**B**) *M. aeruginosa* growth dynamics (n=3, ±SD) of Ma-LMM01-infected (grey line) and non-infected (green line) treatments up to 7 dpi. Created in Biorender.

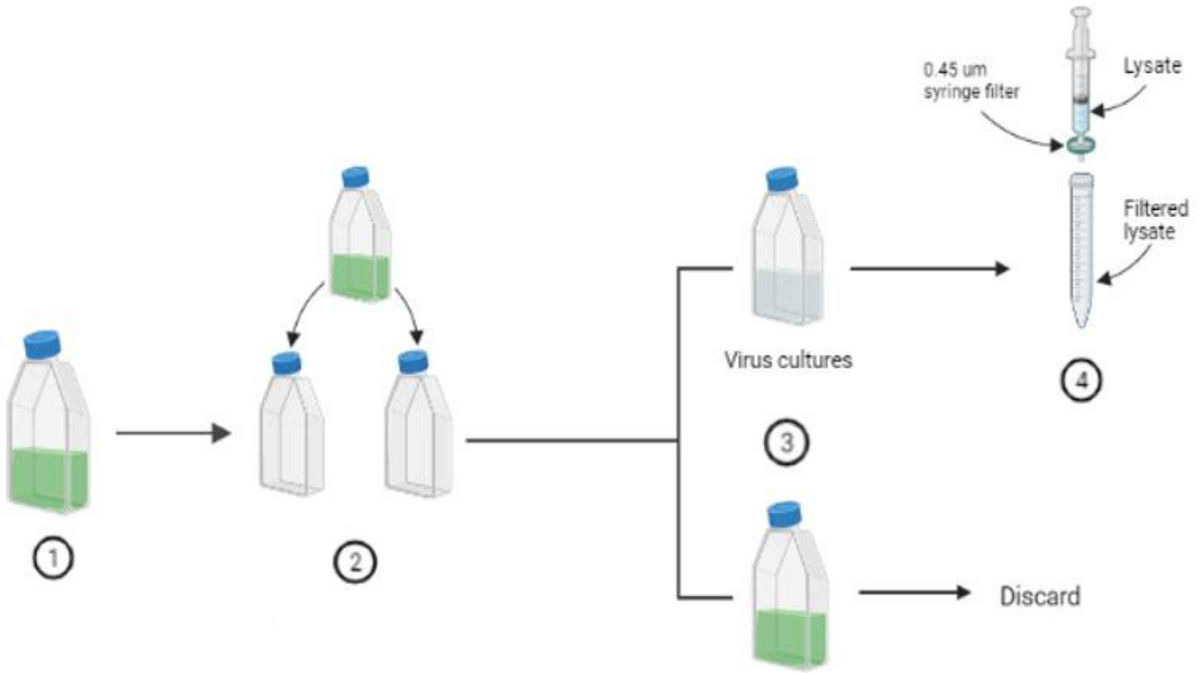


**Figure S2**. Virus propagation on exponentially growing microalgal or cyanobacterial host strains. 1- microalgae and cyanobacteria are incubated in master cultures until mid exponential phase; 2- cultures are split into two; 3- one is infected by a prior stock of a virus lysate and the other serves as a non infected control; 4- infected cultures are filtered though a 0.45 µm filter and the resulting new viral lysate is stored at 4°C in the dark until further use in experiments. Created in Biorender.

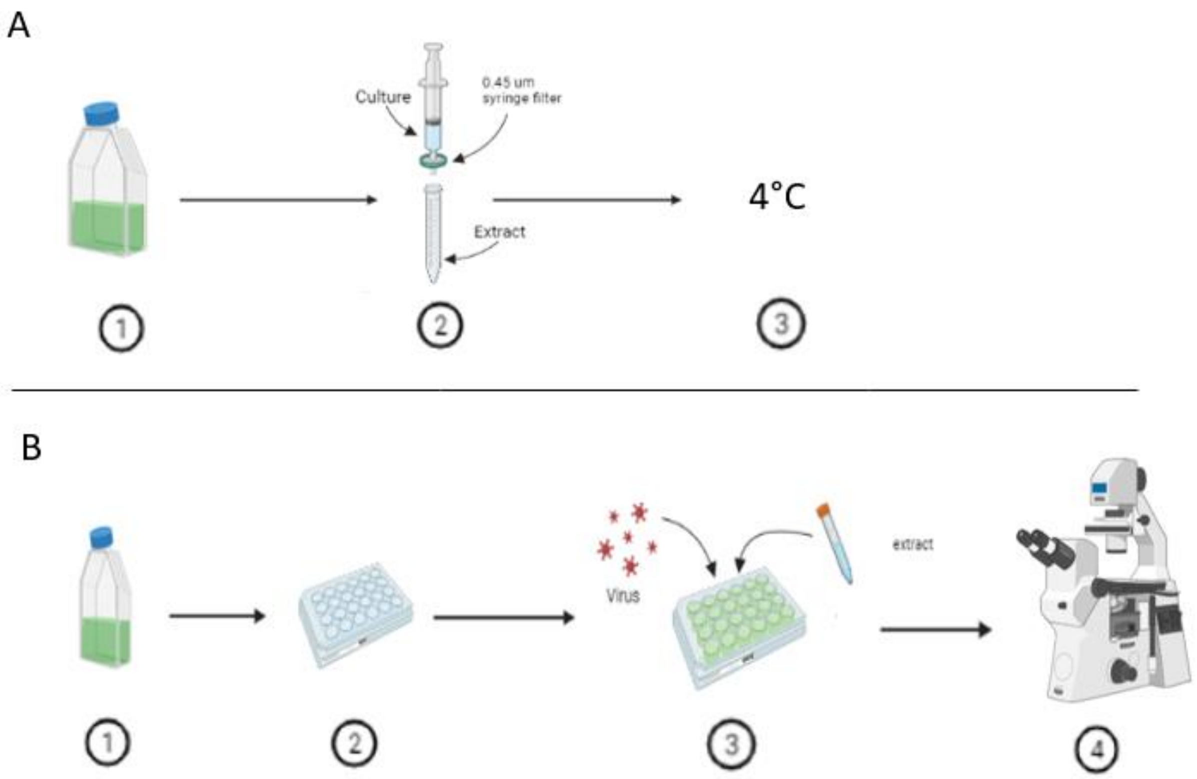


**Figure S3.** Extract filtration from cyanobacteria prior to testing their inhibitory effects against dsDNA, ssDNA and ssRNA viruses. (**A**) 1- cyanobacterial cultures are incubated until late exponential phase; 2- cells are separated from dissolved fractions (“extracts”) by filtering cultures through 0.45 µm filters; 3- filtrates are stored at 4°C until further use in experiments. (**B**) 1- microalgal strains are cultivated until mid-exponential phase; 2- strains are dispensed into 24-well plate as per the outline in Table 2; 3- extracts and viruses are added into the plate as per the outline in Table 2 ; 4- cell abundances are measured by haemocytometery until wells in control plate #3 are cleared. Created in Biorender.

**Table S1**. The viruses and hosts used in our experiments comprise a mixture of strains obtained from colleagues, culture collections, and isolates available in our private culture collection.

| Host | Virus | Virus genome type | Reference | Experiments |
|---|---|---|---|---|
| *M. aeruginosa* NIES-298 | Ma-LMM01 | dsDNA | *(Yoshida et al., 2006)* | Tale 1 |
| *C. tenuissimus* CCAP1010/42 | Cten. DNA virus (CtenDNAV) | ssDNA | *(Tomaru et al., 2011)* | Tale 2 |
| | Cten. RNA virus (CtenRNAV) | ssRNA | *(Shirai et al., 2008)* | |
| *Chlorella variabilis* NC64A | IL-5-2s1 | dsDNA | *(van Etten et al., 1986)* | Tale 2 |
| | NY-2A | | | |
| *Emiliania huxleyi* CCMP374 | E. huxleyi virus-201 (EhV-201) | dsDNA | *(Nissimov et al., 2012)* | Tale 2 |
| | E. huxleyi virus-99B1 (EhV-99B1) | | *(Pagarete et al., 2013)* | |

**Table S2** Sample outline in triplicates for evaluating the antiviral effects of *M. aeruginosa* NIES 298, CPCC 299, and CPCC late exponential growth phase extracts (i.e., filtrates) on CtenDNAV* (V1) and CtenRNAV** (V2) diatom viruses. A= algal host, S= cyanobacterial extract, V= virus. Numbers within wells represent the cyanobacterial strain from which the extract derives. A similar outline was repeated with each algal host-virus pair (e.g., *E. huxleyi*- two EhVs and *C. variabilis*- two chloroviruses).

| **Plate type** | | **A** | **B** | **C** | **D** | **E** | **F** |
|---|---|---|---|---|---|---|---|
| Control plate 1: | **1** | A | | | | | |
| algae only | **2** | A | | | | | |
| | **3** | A | | | | | |
| | **4** | | | | | | |

| | | **A** | **B** | **C** | **D** | **E** | **F** |
|---|---|---|---|---|---|---|---|
| Control plate 2: | **1** | A+298S | | A+299S | | A+300S | |
| algae | **2** | A+298S | | A+299S | | A+300S | |
| + cyanobacterial spike extract | **3** | A+298S | | A+299S | | A+300S | |
| | **4** | | | | | | |

| | | **A** | **B** | **C** | **D** | **E** | **F** |
|---|---|---|---|---|---|---|---|
| Control plate 3: | **1** | A+V1 | | A+V2 | | | |
| algae | **2** | A+V1 | | A+V2 | | | |
| + virus (V1 or V2) | **3** | A+V1 | | A+V2 | | | |
| | **4** | | | | | | |

| | | **A** | **B** | **C** | **D** | **E** | **F** |
|---|---|---|---|---|---|---|---|
| Experimental plate: | **1** | A+298S+V1 | A+299S+V1 | A+300S+V1 | A+298S+V2 | A+299S+V2 | A+300S+V2 |
| algae | **2** | A+298S+V1 | A+299S+V1 | A+300S+V1 | A+298S+V2 | A+299S+V2 | A+300S+V2 |
| + virus (V1 or V2) | **3** | A+298S+V1 | A+299S+V1 | A+300S+V1 | A+298S+V2 | A+299S+V2 | A+300S+V2 |
| + cyanobacterial spike extract | **4** | | | | | | |

* Infection titre of the CtenDNAV virus lysate was $1.59 \times 10^{6}$ viruses $mL^{-1}$.
** Infection titre of the CtenRNAV virus lysate was $2.87 \times 10^{7}$ viruses $mL^{-1}$.
